# Dual-level autoregulation of the *E. coli* DeaD RNA helicase via mRNA stability and Rho-dependent transcription termination

**DOI:** 10.1101/2020.04.19.049098

**Authors:** Sandeep Ojha, Chaitanya Jain

**Affiliations:** Department of Biochemistry and Molecular Biology, University of Miami Miller School of Medicine, Miami, FL 33136, USA.

**Keywords:** RNA helicase, autoregulation, transcription termination, Rho factor

## Abstract

DEAD-box proteins (DBPs) are RNA remodeling factors associated with RNA helicase activity that are found in nearly all organisms. Despite extensive studies on the mechanisms used by DBPs to regulate RNA function, very little is known about how DBPs themselves are regulated. In this work, we have analyzed the expression and regulation of DeaD/CsdA, the largest of the DBPs in *Escherichia coli* (*E. coli*). We show that *deaD* transcription initiates 838 nts upstream of the start of the coding region. We have also found that DeaD is autoregulated through a negative feedback mechanism that operates both at the level of mRNA stability and Rho-dependent transcription termination, and this regulation is dependent upon the 5’ untranslated region (5’ UTR). These findings suggest that DeaD might be regulating the conformation of its own mRNA through its RNA helicase activity to facilitate ribonuclease and Rho access to its 5’UTR.

## INTRODUCTION

DBPs represent a major class of RNA remodeling factors that have been implicated in a variety of RNA regulatory processes in both prokaryotic and eukaryotic organisms (Rocak and Linder 2004; Linder and Jankowsky 2011). The characteristic features of DBPs are a set of >13 conserved motifs distributed over two RecA domains, which jointly bind to RNA and ATP to mediate ATP-dependent RNA or ribonucleoprotein remodeling. *E. coli* contains five members of the DBP family: DbpA, DeaD, RhlB, RhlE and SrmB, which have been implicated in translation regulation, mRNA stability and ribosome biogenesis (Iost et al. 2013; Redder et al. 2015). In particular, an absence of either DeaD or SrmB results in cold-sensitive growth, which is associated with an accumulation of ribosomal precursors (Charollais et al. 2003; Charollais et al. 2004).

In an earlier study, we showed that whereas all five DBPs could be simultaneously removed without loss of viability, the cold-sensitivity of the derivative strain could be largely accounted by the phenotype of the singly mutated *DdeaD* strain (Jagessar and Jain 2010). DeaD is also highly induced during cold shock when cells growing at 37°C are rapidly transferred to 10-16°C, and an absence of this factor impedes the recovery of growth during the acclimatization phase (Jones et al. 1996). Additionally, during cold-shock, DeaD has been found to replace RhlB as the helicase component of the *E. coli* degradosome, an RNA degrading complex, suggesting a role in RNA degradation (Prud’homme-Genereux et al. 2004). Model studies using a chloramphenicol acetyl transferase (*cat*) reporter mRNA showed that DeaD can alleviate the inhibitory effects of secondary structures that overlap the *cat* mRNA ribosome binding site (Butland et al. 2007). Similarly, DeaD has been implicated in the regulation of the *uvrY* mRNA via a translational mechanism (Vakulskas et al. 2014). Additionally, by using crosslinking and immunoprecipitation (CLIP), DeaD was found to cross-link to 39 transcripts *in vivo*, many of which are also regulated by DeaD (Vakulskas et al. 2014). Thus, DeaD appears to have multifaceted roles in gene regulation even though a detailed knowledge of DeaD function in the cell is still incomplete.

Despite the progress made towards defining *E. coli* DBP function, virtually nothing is known about how these factors themselves are regulated. Moreover, any potential regulatory elements in their mRNA 5’ UTR are unknown because their transcription start sites have not been defined or validated. Here we show that *deaD* mRNA transcription starts 838 nts upstream of the coding region, resulting in one of the longest 5’ UTRs known for any *E. coli* transcript. A potential regulatory role for the 5’ UTR is confirmed by the observation that DeaD regulates its own expression, and sequences within the 5’ UTR appear to be important for this process. We show that a high degree of autoregulatory control is mediated by the joint effects of DeaD activity on mRNA stability and transcription elongation. With respect to the latter, the Rho termination factor is shown to play a key role in determining the amount of full-length *deaD* mRNA synthesized inside the cell.

## RESULTS

### The deaD mRNA contains an unusually long 5’ UTR

To study regulatory aspects of *deaD* expression, we first investigated the evidence for *deaD* transcription. A *deaD* mRNA transcription start site (TSS) mapping to 173 nts upstream of the coding region was previously identified by using primer extension, but evidence for such a start site was based on RNA prepared from cells containing a multicopy *deaD* plasmid (Toone et al. 1991). In contrast, high-throughput sequencing studies have shown no evidence for this TSS. Specifically, a recent genome-wide study to map mRNA 5’ ends revealed only three significant TSSs within the first 1000 nts upstream of the *deaD* coding region, of which reads starting 834 nts upstream of the coding region were the most prevalent (Fig. 1A) (Ettwiller et al. 2016). A second study found evidence for only two TSSs starting 834 and 838 nts upstream of the coding region (Thomason et al. 2015). To characterize the *deaD* transcript, northern blot analysis was performed on RNA samples derived from wild-type (WT) and *DdeaD* strains, and a band of ~2.7 kb could be visualized in the former strain (Fig. 1B). With a coding region of 1.89 kb and an expected 3’ UTR of 70 nt, the observed size of the *deaD* mRNA is consistent with transcription initiating ~0.8 kb upstream of the coding region. Interestingly, we noticed the *deaD* transcript runs as a diffuse band on the agarose gels, possibly because of its close migration with the abundant 2.9 kb 23S ribosomal RNA. To define the *deaD* TSS more precisely, we analyzed the *deaD* mRNA by rapid amplification of cDNA ends (RACE), which revealed that the *deaD* TSS starts 838 nt upstream of the coding region, rather than 834 nts (Fig. 1C).

**Fig. 1.**
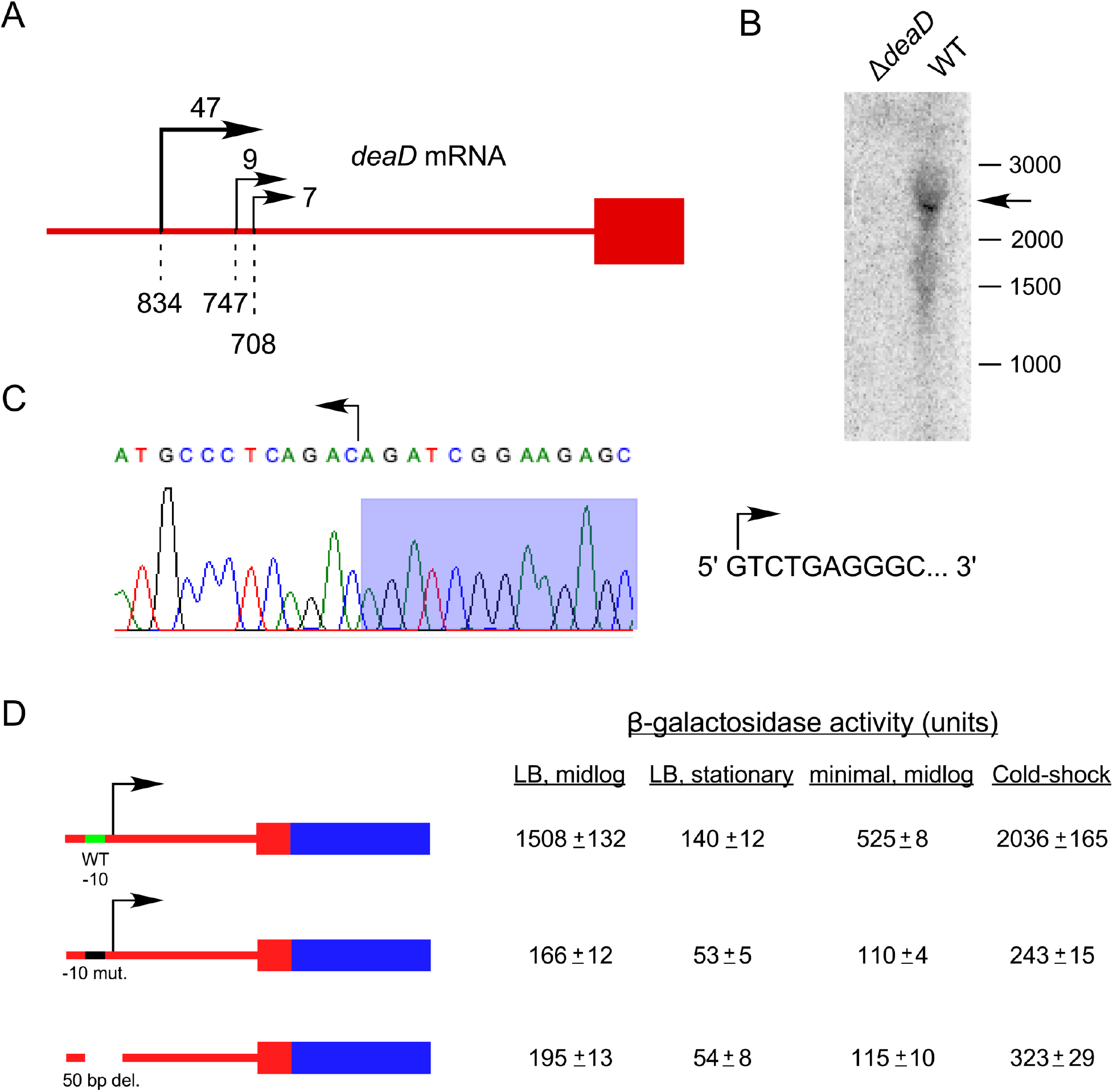
The *deaD* mRNA contains a long 5’ UTR. (A) Schematic description of TSSs upstream of the *deaD* coding region identified by 5’ RNA-Seq. The *deaD* coding region is depicted by a red rectangle and the upstream region by a red line. Transcription start sites identified by Ettwiller et al. (23), are denoted by rightward arrows with relative read scores (RRS) indicated at the top. Only sites with RRS>5 are depicted. Dashed lines and numbers at the bottom indicate the distance between the transcription initiation positions and the start of the coding region. (B) Northern-blot analysis of total RNA isolated from wild type and *DdeaD* strains. The RNA was analyzed using a radiolabeled riboprobe complementary to the *deaD* coding region. The *deaD* mRNA is marked by an arrow. The sizes of molecular weight RNA markers, in nucleotides, run alongside the samples, are shown on the right. (C) TSS determination by RACE. A reverse transcription product generated using total RNA from a WT strain and a primer complementary to the *deaD* 5’ UTR was ligated to a linker, amplified, cloned and sequenced. The resulting trace shows the reverse complement sequence at the junction of *deaD* 5’ UTR and linker sequences (shaded). The sequence of the first ten 5’ nts of the *deaD* mRNA, starting 838 nts upstream of the start codon, is shown at the right. (D) β-galactosidase assays. Plasmids containing a *deaD-lacZ* fusion, a derivative that contains mutations in the putative −10 sequence of the *deaD* promoter, or a 50 bp deletion spanning the −10 and −35 regions, were transformed into a Δ*lacZ* strain and β-galactosidase activity was measured. Regions that correspond to *deaD* and *lacZ* sequences are shown in red and blue, with noncoding and coding regions depicted by lines and rectangles, respectively. β-galactosidase activity was measured after growth at 37°C in LB medium to mid-log phase, in LB medium to stationary phase, in minimal M9 medium to mid-log phase, or after transfer of cultures for one hour to 16°C following an initial growth to mid-log phase in LB medium at 37°C (cold shock).

Next, we were interested in determining the fraction of transcription that originates from the −838 TSS. For these purposes, a plasmid-based *deaD-lacZ* translation fusion and a derivative containing multiple mutations in the putative −10 sequence (TATGAT>CCCCCC) were constructed. Transformation of these plasmids into a *DlacZ* strain resulted in significant β-galactosidase expression for the wild-type construct, but only ~10% residual expression with the mutated construct for cultures grown to midlog phase in LB medium (Fig. 1D), suggesting that nearly 90% of *deaD* mRNA transcription is directed by a promoter that initiates transcription 838 nt upstream of the coding region. Additionally, as most of the activity could be abolished due to mutation, it confirmed that the mapped 5’ end of the *deaD* mRNA corresponds to a genuine TSS, rather than to a processing site. A second construct that contained a deletion spanning from 38 nts upstream of the TSS to 12 nts downstream, including the −35 region of the promoter, yielded a similar decrease, suggesting that the residual transcription in each mutant construct arises from start sites that are not close to the main TSS. To determine the contribution of the −838 TSS to *deaD* transcription under additional conditions, the β-galactosidase activity of the wild-type and mutant *deaD-lacZ* constructs were measured in stationary phase, after growth in minimal medium to mid-log phase or after cold-shock at 16°C. In each case, transcription from the −838 TSS was found to account for most of the signal, indicating that the majority of *deaD* mRNA transcription under different conditions is directed by a single TSS over 800 nts upstream of the coding region.

### The expression of deaD is autoregulated

The presence of a long 5’ UTR in the *deaD* mRNA suggested that DeaD expression could be regulated at multiple levels. One type of regulation, observed for proteins with important physiological cellular roles, is autoregulation, which refers to its ability to buffer its synthesis within a defined range of expression. To determine whether DeaD is autoregulated, mutations within its conserved motifs were introduced into the genomic *deaD* locus. Three different DBP motifs, I, II or VI were targeted, of which motif I and II are also referred to as Walker A and B motifs. Thereafter, RNA was prepared from strains expressing wild-type DeaD, as well as the indicated variants, and analyzed by northern blot (Fig. 2A). Significantly, each mutant strain was found to show elevated levels of *deaD* mRNA. Specifically, expression of the *deaD* mRNA was increased by three-fold in a strain containing a R332Q mutation in motif VI, whereas a higher increase (19-fold) was observed when the first aspartic acid residue in the motif II D-E-A-D sequence was changed to alanine. The further addition of a motif I mutation (K56N) to the D156A mutant increased expression marginally. The K56N + D156A mutant was selected for further analysis and is hereafter referred to as the DM strain.

**Fig. 2.**
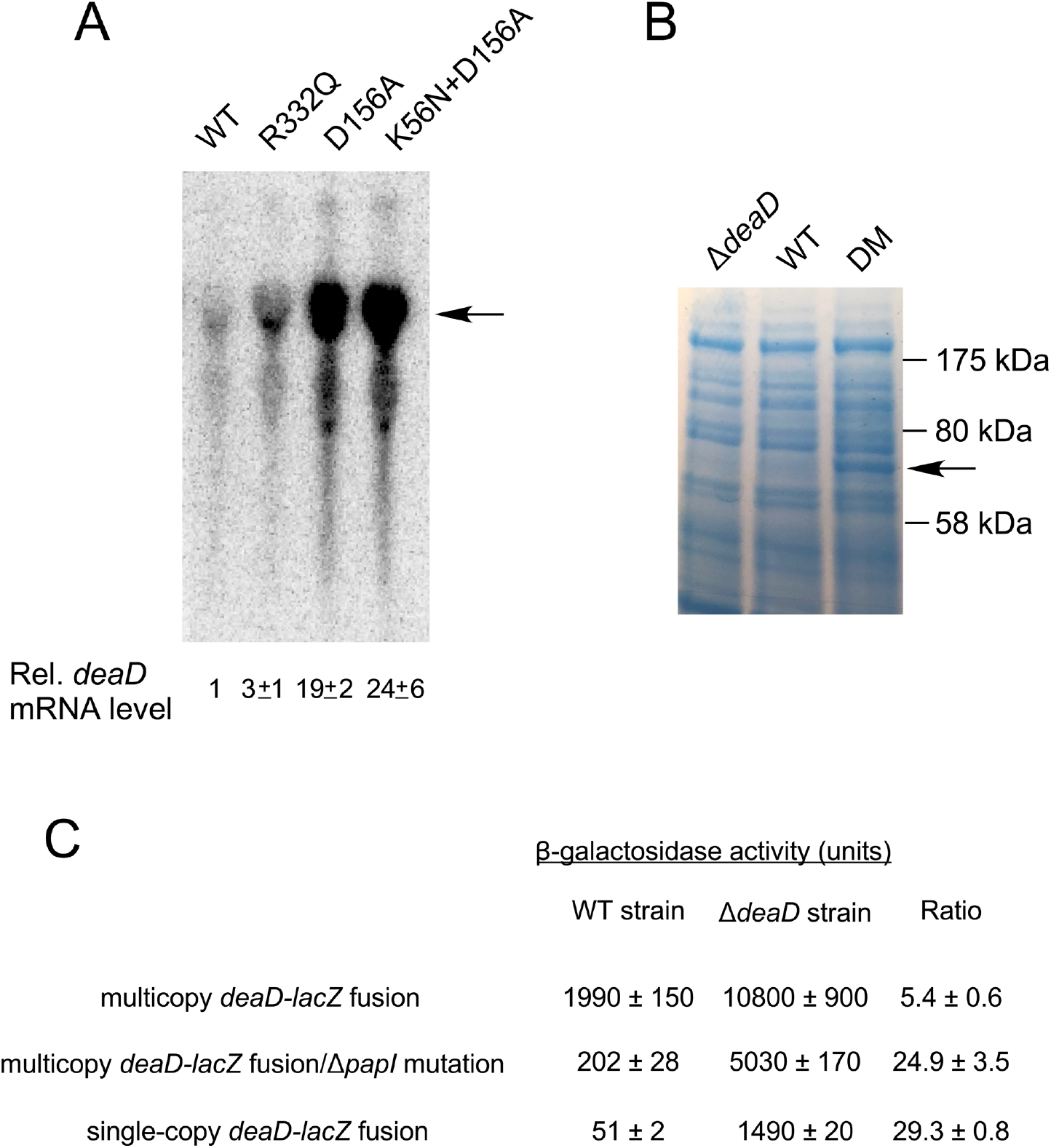
Expression of *deaD* is autoregulated. (A) Expression of *deaD* mRNA in WT strains and strain derivatives containing mutations in conserved motifs of DeaD. The position of the full-length *deaD* mRNA is indicated by an arrow. The normalized amounts of the *deaD* mRNA, along with the associated standard errors, are shown below. (B) Protein analysis. Total cellular proteins were separated by gel electrophoresis and the gels were dye-stained to visualize proteins. The band corresponding to DeaD is indicated by an arrow. The migration of molecular weight markers run alongside the markers is shown on the right. (C) β-galactosidase assays. Assays were performed in WT or Δ*deaD* strains containing multicopy *deaD-lacZ* fusions, in plasmid-containing strains that contained an additional *DpcnB* mutation, or in strains containing a single copy chromosomal *deaD-lacZ* fusion.

To determine whether the increased expression of *deaD* mRNA observed in the DM strain leads to similar increases in protein level, total cellular proteins were directly analyzed by dye staining (Fig. 2B). No DeaD-specific band could be observed for the WT strain in comparison with the *DdeaD* strain, suggesting that the endogenous levels of DeaD are too low to be detected directly. However, in the DM strain, a 70 kilodalton (kDa) protein compatible in size with the 629 amino-acids DeaD protein could be clearly identified, consistent with the significant degree of *deaD* mRNA over-expression observed in the DM strain.

Finally, the multicopy *deaD-lacZ* plasmid was transformed into *lacZ* derivatives of the WT and *DdeaD* strains and β-galactosidase activity was measured (Fig. 2C). Increased expression of the fusion protein was observed in the *DdeaD* strain suggesting that the *deaD* determinants present in the fusion construct, which include the 5’ UTR and the first 100 base-pairs (bps) of the coding region, confer DeaD-dependent regulation. The extent of DeaD-mediated regulation (5-fold) was lower than the increase of mRNA levels in the DM strain (~ 25-fold, Fig. 2A), suggesting that some regulatory regions might be absent in this fusion construct. However, we noticed that the plasmid-transformed *DdeaD* strain exhibited impaired growth, suggesting an alternate possibility that this strain may be unable to express the fusion protein at high levels. To distinguish between these possibilities, we made *DpcnB* derivatives of the WT and *DdeaD* strains to reduce the plasmid copy number, as ColE1 plasmids, such as the one containing the *deaD-lacZ* fusion, propagate at lower copy number in *DpcnB* strains (Xu et al. 2002). As expected, when the plasmid was transformed into the *DpcnB* strain, the amount of *lacZ* expression was reduced by ~10-fold. However, relative to its expression in the *DpcnB* strain, the activity of the *deaD-lacZ* fusion in the Δ*deaD*Δ*pcnB* strain was found to be increased by 25-fold, suggesting that a significantly higher extent of regulation can be achieved when the expression of the fusion is reduced by lowering plasmid copy number. To corroborate these results, the *deaD-lacZ* fusion was transferred to the chromosome in single copy. As was observed with the *DpcnB* strains, the expression of the fusion in the *DdeaD* strain was nearly 30-fold higher than in the WT strain. Based on the inhibited expression of the *deaD-lacZ* fusion under elevated copy number conditions, all downstream experiments were performed using single-copy chromosomal fusions.

Two additional observations are noteworthy. First, the extent to which the activity of the *deaD-lacZ* fusion was increased, when expressed at low or single-copy, was similar to the increase observed for the full-length *deaD* transcript (Fig. 2A), suggesting that the *deaD-lacZ* fusions contain all determinants required for regulation. Second, the similar increase in *deaD* mRNA levels and *deaD-lacZ* translation fusion activity suggested that the observed autoregulation is not likely to involve any significant regulation at the translational level.

### Autoregulation of deaD is jointly mediated by mRNA stability and transcription termination

Although DeaD is implicated in the regulation of several genes, only *cat* and *uvrY* have been studied in some detail (Butland et al. 2007; Vakulskas et al. 2014). In each case, it has been proposed that DeaD regulates these genes through a translational mechanism, possibly by unwinding secondary structures near the translational initiation codon. In theory, DeaD could utilize a similar mechanism for autoregulation, for example, by using its RNA remodeling activity to create secondary structures that impair ribosome binding. However, given that DeaD regulates *deaD* mRNA levels to the same extent as it regulates *deaD-lacZ* translation fusions, it seemed more likely that its autoregulation is effected via mRNA stability or transcription.

To test whether DeaD negatively regulates its expression via mRNA stability, *deaD* mRNA half-life was measured in WT and DM strains. Exponentially growing strains were treated with the RNA polymerase inhibitor, rifampicin, and cells were harvested at various times thereafter for RNA preparation, with an initial time of one min to allow the completion of *deaD* mRNA transcription. The RNA samples were analyzed by northern blot using a *deaD*-specific probe and the amount of mRNA remaining at different times after rifampicin addition was quantified to determine *deaD* mRNA half-life in the WT and mutant strains. Initial efforts to measure half-lives at 37°C did not yield reliable data due to low *deaD* mRNA expression in the WT strain, therefore, the strains were analyzed at 30°C, a condition under which the mRNA levels were found to be increased. These analyses showed that the half-life of the *deaD* mRNA in the WT strain was 2.7 min, but it increased to 7.7 min in the DM strain (Figs. 3A and B). Thus, in the absence of DeaD activity, the *deaD* mRNA is stabilized by nearly three-fold. To identify the enzyme responsible for *deaD* mRNA degradation, the levels of *deaD* mRNA were determined in a WT strain and a derivative containing the temperature-sensitive *rne-3071* mutation. RNase E is implicated in the degradation of most of the *E. coli* mRNAs (Mackie 2013; Mohanty and Kushner 2016), and was thus a likely candidate to be the enzyme that initiates *deaD* mRNA degradation. As expected, the levels of *deaD* mRNA were found to be increased 14-fold in the *rne-3071* strain when cells were harvested after a temperature shift to 42°C (Fig. 3C), implicating RNase E in *deaD* mRNA degradation. In contrast, a smaller difference was seen between the DM and the DM *rne-3071* strain, consistent with the increased stability of the *deaD* mRNA in the former strain.

**Fig. 3.**
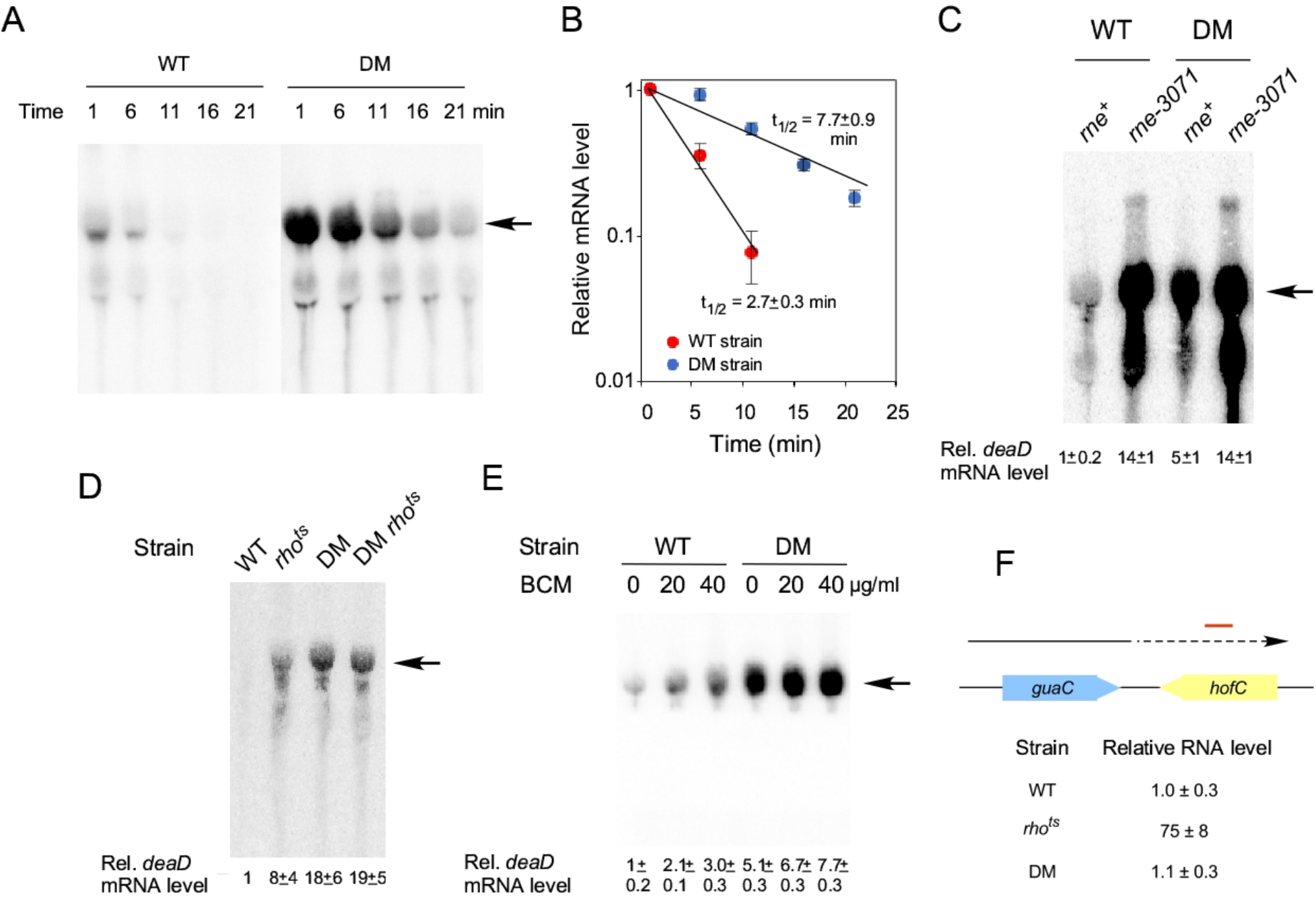
Autoregulation of *deaD* is mediated via mRNA stability and transcription termination. (A) Northern-blot analysis. WT or DM strains were treated with rifampicin and aliquots of the cell cultures were harvested at different times after rifampicin addition, as indicated at the top. Total RNA was prepared from these cultures and analyzed by northern blot using a probe for *deaD* mRNA. (B) Half-life analysis. The levels of *deaD* mRNA from (A) were normalized and plotted as a function of time after rifampicin addition on a semi-logarithmic scale. Each data point represents an average value from three experiments. The calculated half-life of the *deaD* mRNA in each strain is indicated. (C) Inactivation of RNase E increases *deaD* mRNA levels. WT and DM strains, as well as their *rne-3071* derivatives, were transferred to 42°C for 30 min after growth to midlog phase at 30°C, followed by harvesting of cultures, RNA preparation and northern blot analysis. (D) Effects of Rho inactivation on *deaD* mRNA. Total RNA was isolated from WT or DM strains containing either wild-type *rho* or a *rho^ts^* allele. The strains were grown to mid-log phase at 30°C and then transferred to 42°C for 30 min before harvesting. RNA was isolated from the strains and analyzed by northern blotting. (E) Effect of BCM addition on *deaD* mRNA. Cultures of WT or DM strains were grown at 37°C and were either untreated or treated with 20 or 40 μg/ml of BCM for 20 min before harvesting. RNA was isolated from the strains and analyzed by northern blotting. (F) RNA was isolated from WT, *rho^ts^* or DM strains after growth at 42°C for 30 min and the levels of RNA transcription from 538 to 638 bps downstream of the *guaC* coding region were quantified by qRT-PCR. The convergently oriented *guaC* and *hofC* coding regions are shown in blue and yellow, respectively. The arrow depicts transcription from the *guaC* promoter with dotted lines indicating readthrough antisense transcription into the *hofC* coding region. The region analyzed by qRT-PCR is indicated by a red line.

Despite its effects on *deaD* mRNA half-life, the observed three-fold increase in the DM strain was insufficient to explain the ~25-fold increase in steady state levels of the *deaD* transcript (Fig. 2A), suggesting that changes in *deaD* mRNA stability can only partially explain the latter effect. Among the possible explanations for the residual differences were that transcription initiation or elongation could be different between the two strains. However, as DBPs generally act on RNA rather than DNA, any effects of DeaD on the concentration of its own mRNA would be expected to be effected after the transcription initiation stage.

In *E. coli* transcription regulation following initiation commonly occurs either via a factor-independent mode, in which intrinsic RNA sequences cause termination, or by a factor dependent mode, whereby extrinsic factors, most commonly Rho protein, actively dissociates RNA polymerase during transcription (Nudler and Gottesman 2002; Kriner et al. 2016; Ray-Soni et al. 2016). Two approaches were taken to determine whether Rho has a role in *deaD* mRNA regulation. First, we used a strain that contains a temperature sensitive mutation in Rho, an essential protein. Wild-type and *rho^ts^* derivatives of the WT and DM strains were initially grown at a semi-permissive temperature (30°C), then transferred to 42°C, and cells were harvested 30 min after transfer for RNA analysis. Northern-blot analysis of the RNA showed that the amount of fulllength *deaD* mRNA in the WT strain was significantly increased in the *rho^ts^* background, suggesting that Rho also regulates *deaD* mRNA synthesis (Fig. 3D). Additionally, the nearly 20-fold difference in mRNA levels between the WT and the DM strains was reduced to less than three-fold between the *rho^ts^* derivatives of these strains, indicating that regulation of *deaD* mRNA occurs primarily through a Rho-dependent mechanism, with the residual difference attributable to effects on mRNA stability. To further test the role of Rho in *deaD* autoregulation, bicyclomycin (BCM), a Rho inhibitor, was added to growing cultures, and RNA was harvested 20 min after addition of this antibiotic. As with the effects observed with the *rho^ts^* strain, BCM addition was also found to increase *deaD* mRNA levels in the WT strain (Fig. 3E), albeit to a lower degree, possibly because of incomplete Rho inactivation at the BCM concentrations used. To rule out a possibility that *deaD* might be affecting Rho activity indirectly, we used quantitative reverse transcription-PCR (qRT-PCR) to measure the abundance of a validated Rho target. One of the RNA regions whose transcription is sensitive to Rho activity lies downstream of the *guaC* coding region (Dar and Sorek 2018). Transcription over this region is antisense to the convergently oriented downstream *hofC* coding region, and Rho has been shown to be important for suppression of pervasive transcription by antisense transcripts (Peters et al. 2012). As expected, the abundance of the *guaC* mRNA downstream region was dramatically increased upon Rho inactivation (Fig. 3F). In contrast, no such increase was observed in the DM strain, suggesting that the effect of DM mutation on *deaD* mRNA levels is not due to indirect effects on Rho activity. Collectively, these results suggest that DeaD regulates its own mRNA directly through both mRNA degradation and Rho-dependent transcription termination mechanisms.

### Rho-dependent transcription termination in vitro

To further define the action of Rho on *deaD* mRNA, *in vitro* transcription reactions were performed. A DNA template that contains the *deaD* promoter and 1.0 kb of downstream region was incubated with *E. coli* RNA polymerase in the presence of varying amounts of purified Rho protein. In the absence of added Rho, most of the transcripts synthesized during multi-round transcription were found to correspond to the full-length runoff transcript (Fig. 4A, lane 1). The addition of Rho, however resulted in multiple shorter transcripts primarily ranging in size from 400 to 900 nts, consistent with a role of Rho in promoting termination within the *deaD* mRNA (lanes 2-4), whereas the further addition of BCM completely abrogated Rho activity (lane 5). To compare the efficacy of Rho dependent termination on *deaD* mRNA, transcription reactions were performed on a template containing the *rho* gene, since Rho also autoregulates its own synthesis through transcription termination within the 5’ UTR of its own mRNA (Matsumoto et al. 1986). A comparison between the *deaD* and *rho* templates revealed that Rho-dependent termination *in vitro* was more efficient on the *deaD* template than on the *rho* template (Figs. 4B and 4C), consistent with the significant degree of regulation seen *in vivo* (Fig. 2).

**Fig. 4.**
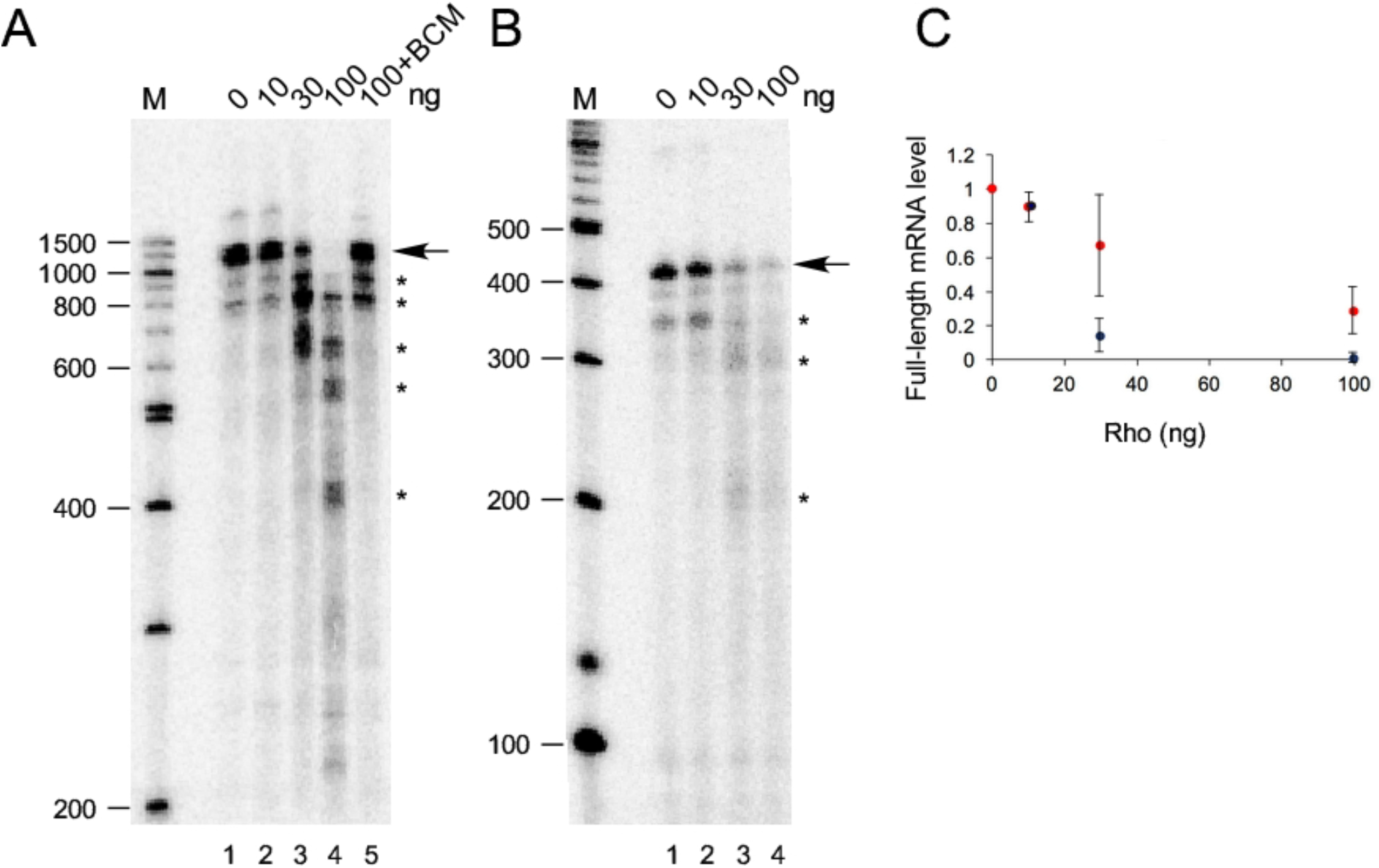
Rho promotes *deaD* mRNA transcription termination *in vitro*. (A) *E. coli* RNA polymerase and different amounts of purified Rho protein, as indicated, were added to a PCR template containing the *deaD* promoter and 955 bps of DNA downstream of the transcription initiation site. The radiolabeled transcription products were resolved by denaturing gel electrophoresis. The fulllength runoff product is indicated by an arrow, whereas prematurely terminated transcripts are indicated by asterisks. Radiolabeled DNA molecular weight markers, run alongside the samples, are shown on the left, along with their sizes in nucleotides. (B) Transcription reactions were performed as in (A), except that a *rho* template was used. (C) Quantitation of transcription termination efficiency using *deaD* and *rho* templates. The relative amounts of full-length mRNA synthesis observed is plotted as a function of Rho concentration. Each data point represents an average from 3-4 replicates. Blue circles, *deaD* template; red circles, *rho* template.

Given that multiple *deaD* mRNA termination products were observed *in vitro*, we investigated whether comparable products could also be seen *in vivo* by northern blot analysis of RNA isolated from WT and DM strains using radiolabeled probes complementary to the 5’ UTR. However, no significant accumulation of short *deaD* mRNA products consistent with Rho-dependent termination could be seen (not shown), suggesting that if such products arise, they might be getting rapidly degraded. Recent transcriptome-wide studies have shown that Rho-dependent termination products often undergo extensive degradation or 3’-end trimming (Dar and Sorek 2018). We also noted that although the results from Fig. 3 indicate that Rho action on the *deaD* mRNA is stimulated by DeaD, prematurely terminated transcripts were observed *in vitro* even without DeaD addition, suggesting a basal level of transcription termination. Efforts to recapitulate enhanced transcription termination by adding purified DeaD to the *in vitro* reactions did not lead to increased termination, either because the conditions used for *deaD* mRNA transcription *in vitro* do not allow DeaD to stimulate Rho termination activity or because DeaD requires other cellular factors to do so *in vivo*.

### Multiple regions in the deaD 5’ UTR contribute to feedback regulation

Given the results obtained implicating DeaD in regulating its own mRNA, we hypothesized that the 5’ UTR plays a key role in mediating this response. To test whether the removal of the 5’ UTR affects the regulation of the *deaD-lacZ* fusion, an initial deletion from 101 to 820 nts upstream of the start codon was made, which upon testing in the WT and *DdeaD* strains was found to exhibit only a 1.4-fold difference between the two strains, in contrast with a 25-fold difference for the wildtype fusion (Fig. 5A). Thus, the 5’ UTR is critical for regulation of the *deaD-lacZ* fusions. Moreover, the loss of regulation as a consequence of an internal deletion distant from the TSS suggests that changes in transcription initiation play little, if any, role in the regulation of *deaD*.

**Fig. 5.**
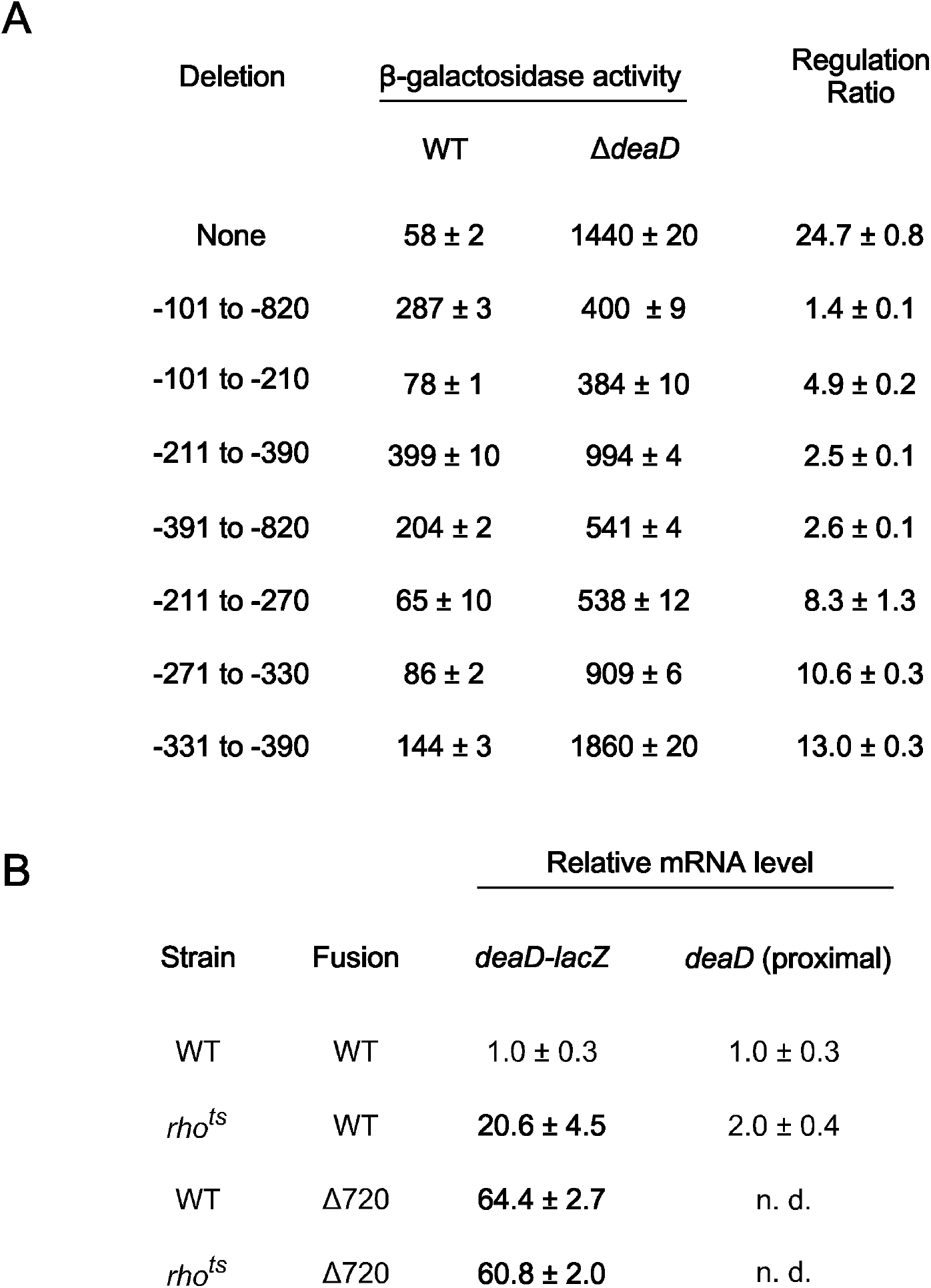
(A) Deletion mapping of regions important for autoregulation. A wild-type single-copy *deaD-lacZ* fusion or variants containing the indicated deletions within the *deaD* 5’ UTR were incorporated into *DlacZ* derivatives of the WT or *DdeaD* strain and the β-galactosidase activity of the resulting strains was measured. (B) RNA was isolated from WT or *rho^ts^* strains containing a WT *deaD-lacZ* fusion or a fusion containing a 720 nt deletion in the *deaD* 5’ UTR. The RNA samples were analyzed by qRT-PCR, with amplification performed either over the junction region of the *deaD-lacZ* transcript (100 nts of *deaD* and 24 nts of *lacZ* sequence) or a 123 nt region near the 5’ end of the transcript. n. d., not determined.

We next tested whether the effect of the 720 nt 5’ UTR deletion can be delimited to a smaller region. Accordingly, we made three smaller deletions that removed 110, 180 or 430 nts of the *deaD* 5’ UTR and tested them both in the WT and *DdeaD* strain backgrounds. Each derivative deletion construct reduced the extent of feedback regulation by five-fold or more relative to the WT fusion, suggesting that they all contribute to the regulation of this transcript, although none of these deletions had as pronounced an effect on regulation as the larger 720 nt deletion.

We further investigated whether the reduced regulatory effect of the 180 nt deletion can be localized to a smaller region, and therefore, we tested three 60 nt deletions spanning this region. Each of the smaller deletions impaired the extent of regulation of the *deaD-lacZ* fusion, though none was as defective as the parental 180 nt deletion. Overall, these results suggest that multiple regions within its large 5’ UTR collectively contribute towards the regulation of the *deaD* mRNA.

As the earlier results had indicated that *deaD* autoregulation is primarily regulated by Rho (Fig. 3D), based on the observation that regulation is lost when the 5’ UTR is deleted, we hypothesized that removal of the 5’ UTR within the context of a *deaD-lacZ* fusion should abolish Rho-dependence. To test that hypothesis, we measured *deaD-lacZ* mRNA levels in both WT and *rho^ts^* strain derivatives containing a WT *deaD-lacZ* fusion by using qRT-PCR. We first performed qRT-PCR using primers that amplify sequences at the *deaD-lacZ* junction, and found that the RNA levels within this region of the transcript were increased by 20-fold in the *rho^ts^* strain, consistent with the effects of Rho-mediated regulation (Fig. 5B). In contrast, when the same analysis was performed on strains containing the 720 nt 5’ UTR deletion, both the WT and *rho^ts^* strains were found to display an equivalent increase in mRNA levels. These results suggest that deletion of the 5’ UTR abrogates Rho-dependent regulation of the *deaD* mRNA. As a control, to verify that the changes observed are not caused by differences in transcription initiation frequency, we also performed qRT-PCR over an initial part of the *deaD* mRNA, 16-138 nts downstream of the TSS. Since Rho generally requires a minimal region to load before it can confer termination (Kriner et al. 2016), it would be expected that any effects of Rho inactivation within the initial regions of a regulated transcript should be reduced. Consistent with this expectation, a relatively smaller two-fold increase in the level of the *deaD* mRNA was observed over the proximal region of this transcript.

## DISCUSSION

DBPs represent a prominent class of RNA-remodeling factors whose functions have been of significant interest to understand how they alter RNA structure to influence gene expression and RNA biology. However, the regulatory aspects of DBP expression have not been investigated in any significant detail. In this work, we have focused on the regulation of DeaD, one of the first DBPs to be identified. An analysis of the *deaD* transcript, as well as the mechanisms by which it is regulated, has led to several novel findings. First, we have found that the *deaD* mRNA contains an 838 nt 5’ UTR, which is longer than the longest primary 5’ UTR (300 nts) annotated in a recent genome-wide study (Thomason et al. 2015). The presence of such an unusually long 5’ UTR suggested that DeaD could be subject to multiple levels of regulation. Indeed, we show that DeaD regulates its own expression, arguably providing the first example of a DBP that is subject to autoregulatory control. Moreover, we have found that *deaD* mRNA is regulated at the level of both mRNA stability and transcription termination. To the best of our knowledge, such a dual level of regulation by an RNA helicase on any transcript has not been reported before. However, examples of dual-level regulation at the both the mRNA stability and transcription termination level by metabolites that bind to riboswitches have been reported for the *lysC* riboswitch in *E. coli* and for the FMN riboswitch that regulates *ribM* expression in *Corynebacterium glutamicum* (Caron et al. 2012; Takemoto et al. 2015; Bastet et al. 2017).

In regard to regulation by transcription termination, we found that the Rho termination factor is important for this process, as Rho mutations or the addition of BCM significantly increased full-length transcript expression (Figs. 3D and 3E). In contrast, strains containing DeaD-inactivating mutations were found to express high levels of *deaD* mRNA even when Rho was functional. The latter observation suggests that DeaD activity is required to remodel *deaD* mRNA so that it becomes accessible to Rho function. This example is reminiscent of control of gene expression by certain riboswitches where ligand dependent changes in the 5’ UTR region enable Rho access, leading to transcription termination (Hollands et al. 2012).

So far, prokaryotic DBPs have been found to regulate gene expression either by regulating translation or RNA stability (Khemici et al. 2005; Resch et al. 2010; Vakulskas et al. 2014). The finding that Rho plays a role in *deaD* regulation thus provides a new mechanism of gene expression regulation by an RNA helicase. We speculate that this mode of regulation is not unique to the *deaD* mRNA and anticipate that future studies will yield additional examples of DeaD, and possibly other DBPs, regulating gene expression through a transcription termination mechanism, rather than via translation or RNA degradation.

To identify the regions that could be important for DeaD autoregulation, we constructed a *deaD-lacZ* translation fusion construct and found that its activity was regulated by DeaD. However, a large 720 nt deletion within the *deaD* mRNA 5’ UTR resulted in an almost complete loss of regulation, indicating that *deaD* regulation is primarily mediated through its 5’ UTR (Fig. 5). We next attempted to determine whether the *cis*-acting elements that control regulation can be localized to a smaller region. Each of three smaller deletions of 110 to 430 nts reduced the extent of regulation, suggesting that the determinants for *deaD* autoregulation are spread out over a large region even though none of these smaller deletions quantitatively captured the effect of the larger deletion.

Based on the results obtained, we suggest that the *deaD* mRNA initially adopts a conformation that contains multiple structured regulatory regions, which upon unwinding by DeaD, become sensitive to Rho and RNase E action. Thus, in the presence of DeaD, *deaD* mRNA undergoes enhanced levels of premature termination, whereas the small fraction of transcripts that escape premature termination get degraded by RNase E. The collective influence of these two processes is a reduction of *deaD* mRNA levels by ~20-fold. This first example of multiple mechanisms used by a DBP to negatively regulate the production of its own transcript underscores the importance of tightly controlling any fluctuations in its expression inside the cell. We expect that future studies will reveal additional examples of autoregulatory control not only for other DBPs but also for RNA-remodeling factors that perform important functions inside the cell.

## MATERIALS AND METHODS

### Strains and plasmids

Strains were derived from MG1655*, a derivative of MG1655 that contains a point mutation in *rph* that restores the reading frame of its prematurely terminated gene product (Gutgsell and Jain 2012). Δ*deaD*, Δ*lacZ* and Δ*pcnB* mutations were derived from the Keio collection of strains containing the respective deletion alleles marked with a kanamycin-resistant (*kan^R^*) element, which were transferred to MG1655* by P1 transduction (Baba et al. 2006). For the *DdeaD::kan* allele, the *kan^R^* marker was removed by using FLP recombinase. Point mutations in *deaD* were introduced via two steps. First, *deaD* codons 48-339 were replaced by a *tetA-sacB* cassette by using recombineering (Li et al. 2013). Second, PCR products spanning the deleted *deaD* region and containing the desired mutations were recombined into the derivative strain and grown on plates containing fusaric acid and sucrose, which provide a counter-selection against the *tetA-sacB* cassette. Colonies growing under the selection conditions were isolated and sequenced to confirm the presence of the desired mutations. A strain containing a *rho^ts^* allele was obtained from Dr. Sankar Adhya, and the *rho^ts^* allele was transduced into MG1655* by P1 transduction using a linked *ilvC::kan* marker (Gulletta et al. 1983). An *rne^ts^* allele (*rne-3071*) was derived from strain CJ1826 (Jain and Belasco 1995) and transduced to recipient strains using a linked *plsX::*chloramphenicol-resistant marker.

The *deaD-lacZ* plasmid was constructed by cloning PCR amplified *deaD* DNA spanning from 973 bp upstream of the coding region to 100 bp downstream between the BamHI and XhoI sites of pLACZY1A, a ColE1-based plasmid with expected copy number of ~20/cell (Jain 1993). Derivatives of this plasmid were made using a Q5 site-directed mutagenesis kit (NEB). Single copy *deaD-lacZ* fusions were made by recombining the plasmid-borne fusions with modified *λ* phage, integrating the recombinant phage at the *λ att* site on the chromosome, and identifying monolysogens by PCR-based screening (Simons et al. 1987; Powell et al. 1994).

### Northern blotting

Total RNA was isolated using the hot-phenol method. RNA samples were fractionated on 1.2% agarose gels in MOPS buffer containing 0.66 M formaldehyde. After electrophoresis, the RNA was transferred to positively charged membrane (Nytran), cross-linked using 254 nM ultraviolet light, and hybridized overnight with radiolabeled riboprobes at 65°C. The DNA template for riboprobe synthesis was made by PCR using *E. coli* DNA as a template and primers with the sequences 5’-AATTTAATACGACTCACTATAGGTTACGCATCACCACCGAAAC-3’ and 5’- GTTCGTCATATCGTTGGTGC-3’. Riboprobe synthesis was performed using the HiScribe kit (NEB) with γ-^32^P UTP added. For mRNA half-life measurements, samples of total cellular RNA were isolated periodically beginning 1 min after rifampicin addition to a final concentration of 200 μg/ml. All experiments were performed a minimum of three times.

### In vitro transcription termination assays

Transcription reactions were performed on a 1.1 kb DNA template synthesized by using PCR on *E. coli* DNA. The sequences for the primers used to synthesize the *deaD* template were 5’-CGGTACCATTGCAACCGACTTTAC-3’ and 5’-ATACACTCTGCCTGAATTGG 3’. The primer sequences for synthesis of a 0.4 kb *rho* template were 5’-TGGACGCCCGGCGTGAGTCA-3’ and 5’-GCAAAAATAATGTCCTGCTT-3’. Transcription reactions, performed in 10 μl, contained 10 ng of template in 10% v/v glycerol, 40 mM Tris–HCl (pH 7.9), 50 mM KCl, 5mM MgCl2, 0.1mM dithiothreitol, 1 mM ATP, GTP and CTP, 20 μM UTP, 1μCi γ-^32^P UTP (3000 Ci/mmol) and 1 unit *E. coli* RNA polymerase holoenzyme (NEB). The reactions were supplemented with Rho and BCM (100 μM), as indicated. After incubation for 20 min at 37°C, the reactions were terminated by adding 2 μl of acidic phenol, and the samples were vortexed and centrifuged. 7 μl of the aqueous phase was removed and added to 5 μl of loading dye. The samples were heated at 95°C for 5 min and the transcription products were resolved on denaturing 4% polyacrylamide gel and visualized using a Personal Molecular Imager (BioRad).

### 5’ RACE

Primer extension was performed on RNA from a WT strain as described (Diwa et al. 2000), using a primer with the sequence 5’-CAGAATCAAACGCTTCATAG-3’. The cDNA product was precipitated and 1 μl of the cDNA product was added to 1 μl of a 5’ phosphorylated, 3’ blocked oligonucleotide (5’-AGATCGGAAGAGCACACGTCT-NH2-3’) at 1 μM and 1 μl of dimethyl sulfoxide. The mix was heated to 75°C for three min and placed on ice. The reaction was supplemented with 3 μl of T4 RNA ligase I (NEB), 1.5 μl of 10X reaction buffer, 6.5 μl of 50% PEG-8000 and 1 μl of 10mM ATP. The reaction was incubated at room temperature for one hour and an aliquot was used to amplify the ligation product using primers with the sequences 5’- CCTAAGTAATTGAATACTTC-3’ and 5’-AGACGTGTGCTCTTCCGATCT-3’. The PCR product was cloned into pGEM-T easy (Promega), amplified by using primers with the sequences 5’- AGTCACGACGTTGTAAAACG-3’ and 5’-GAGCGGATAACAATTTCACACAGG-3’, and the resulting product was sequenced using Sanger sequencing (Genewiz).

### qRT-PCR

One μg of total RNA was treated with 0.4 units of DNase I (Promega) and 0.2 units of Turbo-DNase (Ambion) in 1X DNase buffer for 45 min at 37°C in a volume of 10 μl. 90 μl of water was added to these reactions and 2 μl of each mix was used for qRT-PCR using the Luna qRT-PCR kit from New England Biolabs (cat # E3005) and 200 nM of primers in a total volume of 5 μl using a BioRad CFX 96 instrument. The sequences of the primers used for quantification of RNA downstream of *guaC* mRNA were 5’-GCAACTTATTGGCTATCGCC-3’ and 5’- GGCACTAACGCAGGGGATC-3’; for quantification of *deaD* mRNA were 5’- CGCGTAACGATTTTTCGCAAGCG-3’ and 5’-CAGAATCAAACGCTTCATAG-3’; and for quantification of the *deaD-lacZ* fusion RNA were 5’-GATGGCTGAATTCGAAA-3’ and 5’- AGTCACGACGTTGTAAAACG-3’. ΔC_t_ measurements were converted into fold-differences assuming an amplification efficiency of two per PCR cycle. All RNA quantitation measurements reported are based on 3-4 replicates, with standard errors indicated.

### Protein analysis

Cell extracts were prepared by heating cells resuspended in sample buffer at 100°C for 10 min. The cells were fractionated on a 4-15% gel (BioRad) and the gels were stained using PageBlue protein staining solution (Fermentas).

### β-galactosidase assays

β-galactosidase assays were performed as described (Jain and Belasco 2000). The β-galactosidase activities from *deaD-lacZ* plasmids or single-copy fusions were measured in Δ*lacZ* derivatives of the WT and Δ*deaD* strains. The β-galactosidase activity of endogenous *lacZ* was measured using strain MG1655* in LB medium supplemented with 1 mM IPTG. All β-galactosidase values reported are based on 3-7 replicates, with standard errors indicated.

## ACKNOWLEDGEMENTS

We thank Dr. Evgeny Nudler (New York University) for the gift of purified Rho protein. This work was supported by Grant GM114540 from the National Institutes of Health.

## REFERENCES

Baba T, Ara T, Hasegawa M, Takai Y, Okumura Y, Baba M, Datsenko KA, Tomita M, Wanner BL, Mori H. 2006. Construction of Escherichia coli K-12 in-frame, single-gene knockout mutants: the Keio collection. Mol Syst Biol 2: 2006 0008.

Bastet L, Chauvier A, Singh N, Lussier A, Lamontagne AM, Prevost K, Masse E, Wade JT, Lafontaine DA. 2017. Translational control and Rho-dependent transcription termination are intimately linked in riboswitch regulation. Nucleic acids research 45: 7474–7486.

Butland G, Krogan NJ, Xu J, Yang WH, Aoki H, Li JS, Krogan N, Menendez J, Cagney G, Kiani GC et al. 2007. Investigating the in vivo activity of the DeaD protein using protein-protein interactions and the translational activity of structured chloramphenicol acetyltransferase mRNAs. J Cell Biochem 100: 642–652.

Caron MP, Bastet L, Lussier A, Simoneau-Roy M, Masse E, Lafontaine DA. 2012. Dual-acting riboswitch control of translation initiation and mRNA decay. Proceedings of the National Academy of Sciences of the United States of America 109: E3444–3453.

Charollais J, Dreyfus M, Iost I. 2004. CsdA, a cold-shock RNA helicase from Escherichia coli, is involved in the biogenesis of 50S ribosomal subunit. Nucleic acids research 32: 2751–2759.

Charollais J, Pflieger D, Vinh J, Dreyfus M, Iost I. 2003. The DEAD-box RNA helicase SrmB is involved in the assembly of 50S ribosomal subunits in Escherichia coli. Molecular microbiology 48: 1253–1265.

Dar D, Sorek R. 2018. High-resolution RNA 3’-ends mapping of bacterial Rho-dependent transcripts. Nucleic acids research 46: 6797–6805.

Diwa A, Bricker AL, Jain C, Belasco JG. 2000. An evolutionarily conserved RNA stem-loop functions as a sensor that directs feedback regulation of RNase E gene expression. Genes Dev 14: 1249–1260.

Ettwiller L, Buswell J, Yigit E, Schildkraut I. 2016. A novel enrichment strategy reveals unprecedented number of novel transcription start sites at single base resolution in a model prokaryote and the gut microbiome. BMC Genomics 17: 199.

Gulletta E, Das A, Adhya S. 1983. The pleiotropic ts15 mutation of E. coli is an IS1 insertion in the rho structural gene. Genetics 105: 265–280.

Gutgsell NS, Jain C. 2012. Gateway role for rRNA precursors in ribosome assembly. Journal of bacteriology 194: 6875–6882.

Hollands K, Proshkin S, Sklyarova S, Epshtein V, Mironov A, Nudler E, Groisman EA. 2012. Riboswitch control of Rho-dependent transcription termination. Proceedings of the National Academy of Sciences of the United States of America 109: 5376–5381.

Iost I, Bizebard T, Dreyfus M. 2013. Functions of DEAD-box proteins in bacteria: current knowledge and pending questions. Biochim Biophys Acta 1829: 866–877.

Jagessar KL, Jain C. 2010. Functional and molecular analysis of Escherichia coli strains lacking multiple DEAD-box helicases. RNA (New York, NY) 16: 1386–1392.

Jain C. 1993. New improved lacZ gene fusion vectors. Gene 133: 99–102.

Jain C, Belasco JG. 1995. RNase E autoregulates its synthesis by controlling the degradation rate of its own mRNA in Escherichia coli: unusual sensitivity of the rne transcript to RNase E activity. Genes Dev 9: 84–96.

Jain C, Belasco JG. 2000. Rapid genetic analysis of RNA-protein interactions by translational repression in Escherichia coli. Methods Enzymol 318: 309–332.

Jones PG, Mitta M, Kim Y, Jiang W, Inouye M. 1996. Cold shock induces a major ribosomal-associated protein that unwinds double-stranded RNA in Escherichia coli. Proceedings of the National Academy of Sciences of the United States of America 93: 76–80.

Khemici V, Poljak L, Toesca I, Carpousis AJ. 2005. Evidence in vivo that the DEAD-box RNA helicase RhlB facilitates the degradation of ribosome-free mRNA by RNase E. Proceedings of the National Academy of Sciences of the United States of America 102: 6913–6918.

Kriner MA, Sevostyanova A, Groisman EA. 2016. Learning from the Leaders: Gene Regulation by the Transcription Termination Factor Rho. Trends Biochem Sci 41: 690–699.

Li XT, Thomason LC, Sawitzke JA, Costantino N, Court DL. 2013. Positive and negative selection using the tetA-sacB cassette: recombineering and P1 transduction in Escherichia coli. Nucleic acids research 41: e204.

Linder P, Jankowsky E. 2011. From unwinding to clamping - the DEAD box RNA helicase family. Nat Rev Mol Cell Biol 12: 505–516.

Mackie GA. 2013. RNase E: at the interface of bacterial RNA processing and decay. Nat Rev Microbiol 11: 45–57.

Matsumoto Y, Shigesada K, Hirano M, Imai M. 1986. Autogenous regulation of the gene for transcription termination factor rho in Escherichia coli: localization and function of its attenuators. Journal of bacteriology 166: 945–958.

Mohanty BK, Kushner SR. 2016. Regulation of mRNA Decay in Bacteria. Annu Rev Microbiol 70: 25–44.

Nudler E, Gottesman ME. 2002. Transcription termination and anti-termination in E. coli. Genes Cells 7: 755–768.

Peters JM, Mooney RA, Grass JA, Jessen ED, Tran F, Landick R. 2012. Rho and NusG suppress pervasive antisense transcription in Escherichia coli. Genes Dev 26: 2621–2633.

Powell BS, Rivas MP, Court DL, Nakamura Y, Turnbough CL, Jr. 1994. Rapid confirmation of single copy lambda prophage integration by PCR. Nucleic acids research 22: 5765–5766.

Prud’homme-Genereux A, Beran RK, Iost I, Ramey CS, Mackie GA, Simons RW. 2004. Physical and functional interactions among RNase E, polynucleotide phosphorylase and the coldshock protein, CsdA: evidence for a ‘cold shock degradosome’. Molecular microbiology 54: 1409–1421.

Ray-Soni A, Bellecourt MJ, Landick R. 2016. Mechanisms of Bacterial Transcription Termination: All Good Things Must End. Annual review of biochemistry 85: 319–347.

Redder P, Hausmann S, Khemici V, Yasrebi H, Linder P. 2015. Bacterial versatility requires DEADbox RNA helicases. FEMS Microbiol Rev 39: 392–412.

Resch A, Vecerek B, Palavra K, Blasi U. 2010. Requirement of the CsdA DEAD-box helicase for low temperature riboregulation of rpoS mRNA. RNA Biol 7: 796–802.

Rocak S, Linder P. 2004. DEAD-box proteins: the driving forces behind RNA metabolism. Nat Rev Mol Cell Biol 5: 232–241.

Simons RW, Houman F, Kleckner N. 1987. Improved single and multicopy lac-based cloning vectors for protein and operon fusions. Gene 53: 85–96.

Takemoto N, Tanaka Y, Inui M. 2015. Rho and RNase play a central role in FMN riboswitch regulation in Corynebacterium glutamicum. Nucleic acids research 43: 520–529.

Thomason MK, Bischler T, Eisenbart SK, Forstner KU, Zhang A, Herbig A, Nieselt K, Sharma CM, Storz G. 2015. Global transcriptional start site mapping using differential RNA sequencing reveals novel antisense RNAs in Escherichia coli. Journal of bacteriology 197: 18–28.

Toone WM, Rudd KE, Friesen JD. 1991. deaD, a new Escherichia coli gene encoding a presumed ATP-dependent RNA helicase, can suppress a mutation in rpsB, the gene encoding ribosomal protein S2. Journal of bacteriology 173: 3291–3302.

Vakulskas CA, Pannuri A, Cortes-Selva D, Zere TR, Ahmer BM, Babitzke P, Romeo T. 2014. Global effects of the DEAD-box RNA helicase DeaD (CsdA) on gene expression over a broad range of temperatures. Molecular microbiology 92: 945–958.

Xu FF, Gaggero C, Cohen SN. 2002. Polyadenylation can regulate ColE1 type plasmid copy number independently of any effect on RNAI decay by decreasing the interaction of antisense RNAI with its RNAII target. Plasmid 48: 49–58.

